# A functional logic for neurotransmitter co-release in the cholinergic forebrain pathway

**DOI:** 10.1101/2021.02.25.432623

**Authors:** Aditya Nair, Martin Graf, Yue Yang Teo, George J. Augustine

**Author notes:** Computation and Neural Systems, California Institute of Technology, Pasadena, CA, USA. Corresponding author: George Augustine.

## Abstract

The forebrain cholinergic system has recently been shown to co-release both acetylcholine and GABA. We have discovered that such co-release by cholinergic inputs to the claustrum differentially affects neurons that project to cortical versus subcortical targets. The resulting changes in neuronal gain toggles network efficiency and discriminability of output between two different projection subcircuits. Our results provide a potential logic for neurotransmitter co-release in cholinergic systems.

## Main Text

The cholinergic system of the basal forebrain is a crucial pathway that modulates attention, arousal and learning^1–3^. Such actions arise from the ability of the cholinergic system to alter neuronal excitability and shape the correlational structure of neural populations^4–6^. The prevailing view that the cholinergic system implements such computations solely by releasing acetylcholine (ACh) has been challenged by the recent discovery that nearly all forebrain cholinergic neurons co-release the inhibitory transmitter, GABA, along with ACh^7,8^. While such co-release has been observed in multiple brain areas, a functional logic for it is missing: does co-release happen in a target-specific manner and how does it impact cholinergic computations?

We have addressed these questions by analyzing cholinergic modulation of the claustrum. The claustrum receives input from basal forebrain cholinergic neurons^11–13^ and has been implicated in attention, perhaps by altering cortical gain^11,14^. Cholinergic modulation of claustrum neurons was examined by whole-cell patch clamp recordings in brain slices from a ChAT-Cre mouse line crossed with another line with Cre-dependent expression of ChR2-YFP^15^, thereby targeting ChR2 exclusively to cholinergic neurons. To isolate cholinergic responses, recordings were performed in the presence of a glutamate receptor blocker (kynurenic acid, KYN; 1 μM) and GABA receptor blocker (Gabazine, GBZ; 10 μM). The claustrum consists of multiple types of projection neurons and interneurons^16^. A trained classifier was applied to whole-cell patch clamp measurements of intrinsic electrical properties to identify neurons that project to cortical or non-cortical targets, as well as to identify the three known types of local interneurons^16^.

ChR2-mediated photostimulation of cholinergic input elicited responses in claustro-cortical (CC), claustro-subcortical (CS) projection neurons and VIP interneurons (VIP-IN; Fig 1a). Only a small fraction (5%, n = 5/97 neurons) of CC neurons received cholinergic excitatory input, while five times more CS neurons (25%, n = 16/64 neurons) and even more VIP-IN (44%, n = 4/9 neurons) received such excitation (Fig 1b, EPSC amplitude: CC: 14 ± 1.2 pA, CS: 35.3 ± 5 pA, VIP: 64.7 ± 5.5 pA). These were monosynaptic inputs, because they persisted after tetrodotoxin (TTX; 1 μM) was used to block action potentials and 4-AP was applied to enhance ChR2-mediated depolarization^17–19^ (Fig 1c). These excitatory responses were mediated by nicotinic ACh receptors, because they were blocked by a nicotinic receptor blocker (mecamylamine, MECA; 10 μM; Fig 1c) and had reversal potentials near zero (Fig S1a), typical of responses mediated by nicotinic receptors^20^. SST and PV interneurons never responded to cholinergic photostimulation (Fig 1b). These results reveal cell-type specific modulation of claustrum neurons and are consistent with reports that VIP interneurons are a critical target for cholinergic modulation in other parts of the brain^21,22^.

**Figure 1:**
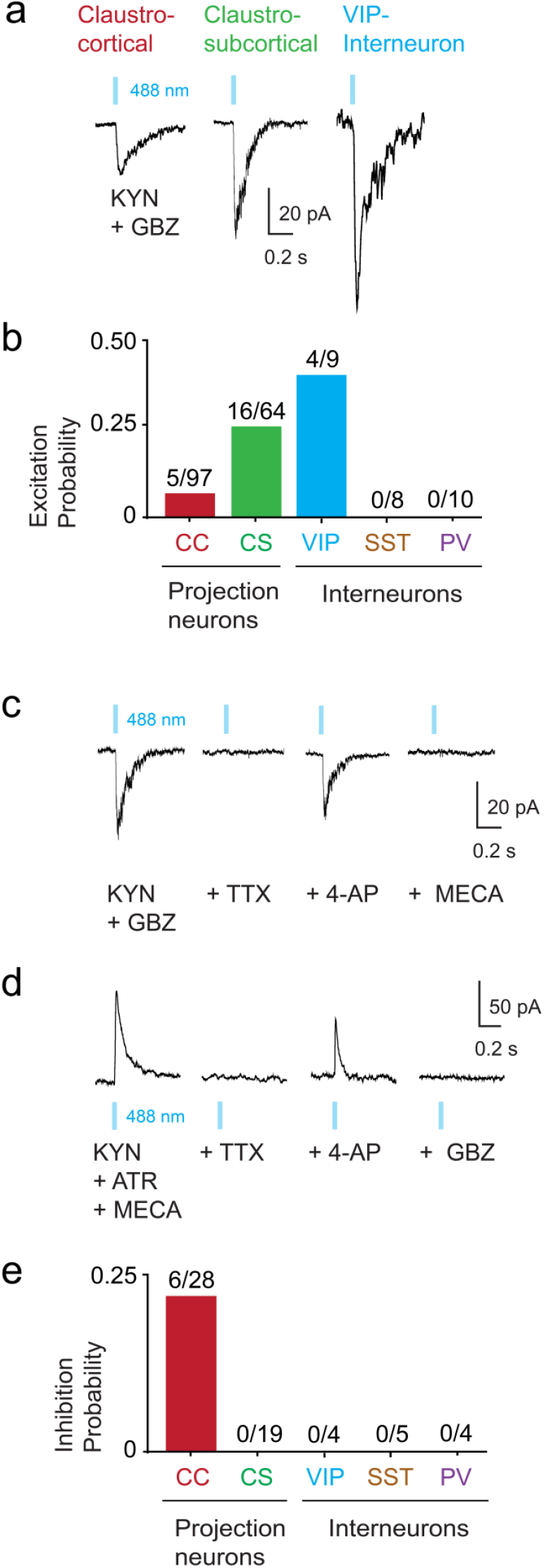
Cell-type specific direct cholinergic inputs in claustrum neurons. **a.** Direct excitatory inputs to claustrum neurons measured in the presence of Kynurenic Acid (KYN, 1 μM) and Gabazine (GBZ, 10 μM) by photoactivating cholinergic terminals using 50ms light pulses at 488 nm. **b.** Distribution of excitation probability in all cell-types tested for direct excitation Cells held at −40 mV. **c.** A subset of neurons were tested for monosynaptic connectivity using Tetrodotoxin (TTX, 1 μM) and 4- Aminopyridine (4-AP, 500 μM). **d.** CC neurons receive direct inhibition measured using KYNA, Atropine (ATR) and Mecamylamine (MECA). Monosynaptic connectivity is tested using TTX and 4-AP. **e**. Distribution of inhibition probability in all cell-types tested for direct inhibition.

Because the inhibitory neurotransmitter, GABA, can be co-released by cholinergic neurons^7^, we asked whether claustrum neurons receive GABAergic inhibition by photostimulating cholinergic neurons after blocking excitatory responses with KYN, MECA and atropine (ATR; 10 μM). Under such conditions, monosynaptic outward currents were observed (Fig 1d, IPSC amplitude: 49.3 ± 6.2 pA), that persisted in the in the presence of TTX and 4-AP. These responses were mediated by GABA_A_ receptors, because they were blocked by GBZ (Fig 1d) and were inhibitory because their reversal potential of −70 mV was more negative than action potential threshold of CC neurons (−33.8 ± 0.2 mV, Fig S1b). Remarkably, we only found inhibitory inputs to CC neurons (Fig 1e). Indeed, CC neurons were much more likely to exhibit inhibitory responses to cholinergic input (21%, n = 6/28 neurons) than excitatory responses (5%). These results reveal a logic for cholinergic co-release of GABA: while claustral neurons projecting to subcortical structures, as well as VIP-IN, are excited via cholinergic activation of nicotinic receptors, neurons projecting to cortical structures are more likely to be inhibited by co-released GABA. Such opposing regulation by cholinergic input may also be present in cortical circuits^23^.

To understand the functional consequences of the dual modulation produced by co-transmitter release, we determined the effects of cholinergic input on action potentials (APs) evoked by depolarizing current pulses using a 1-second long train of blue light (488 nm) pulses delivered at 10Hz. In CC neurons, cholinergic input reduced the frequency of APs evoked by a 100 pA depolarizing current pulse (Fig 2a, left, n = 12 neurons), presumably due to the inhibitory action of GABA. In contrast, this input increased AP firing in both CS neurons and VIP neurons, presumably due to ACh excitation (Fig 2b,c, left, n = 10 CS neurons, 5 VIP-IN). The slope of the relationship between current magnitude and AP frequency, the input-output (IO) curve, reveals neuronal gain^24^; changes in gain are a characteristic feature of cholinergic modulation of the cortex^25^. While inhibitory input decreased the gain of CC neurons (Fig 2a, right), excitatory input increased the gain of CS neurons and VIP-IN (Fig 2b,c, right). Increased modulation of CS neuron gain was also accompanied by a secondary decrease in AP frequency at higher current intensities (Fig 2b, right). Thus, the co-transmitters released by cholinergic input produces opposing, cell-type specific gain modulation of claustrum neurons (Fig 2d).

**Figure 2:**
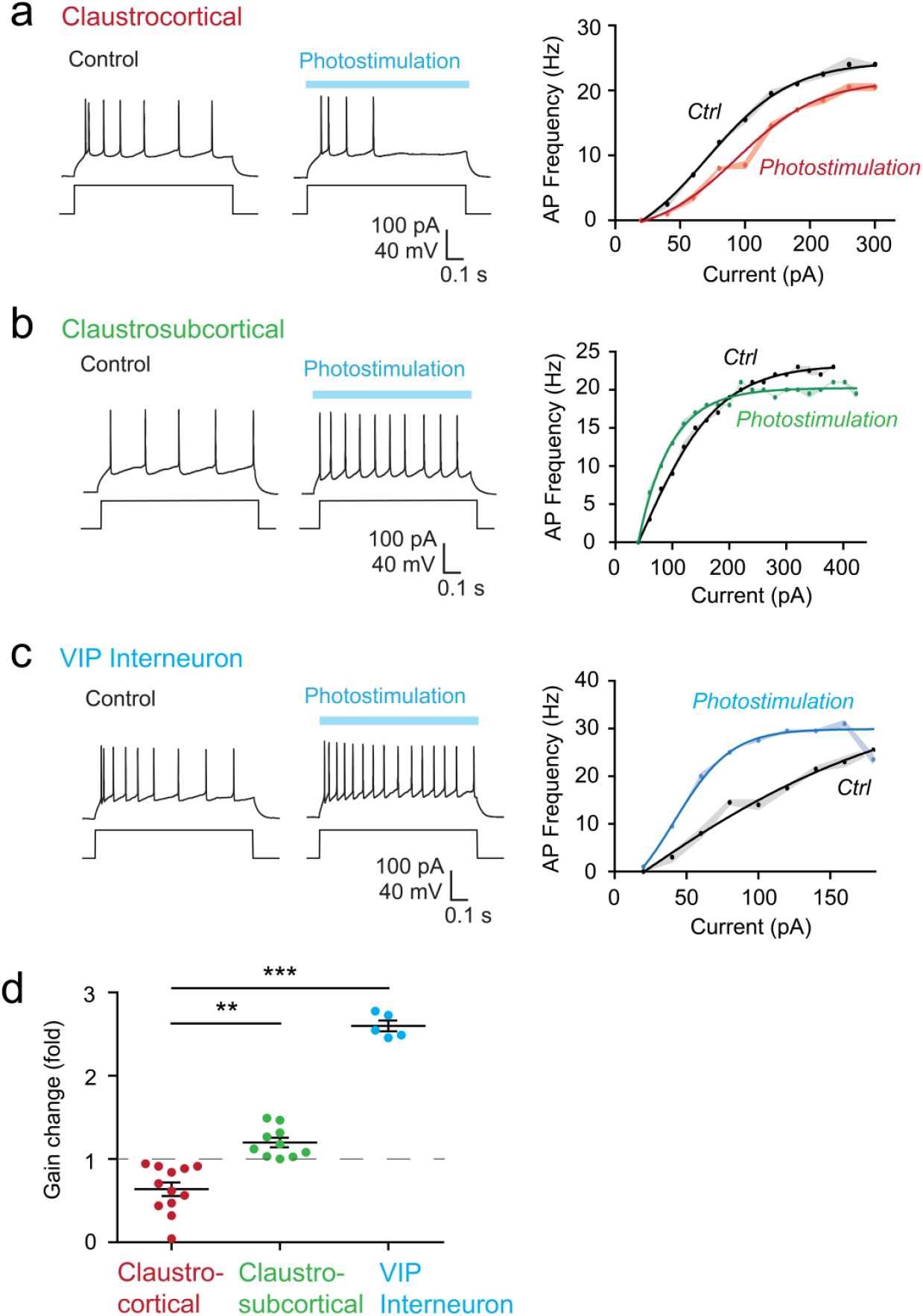
Opposing gain control of claustrum cell-types. Left: Responses of claustrum neurons to 100 pA current injection during control and cholinergic photostimulation: 20ms pulses at 10 Hz for one second (**a**: Claustrocortical, **b**: Claustrosubcortical, **c**: Putative VIP interneurons). Right: Quantification of changes on input-current output-frequency (IO) curve in control and cholinergic photostimulation conditions. **d**. Quantification of changes on neuronal gain (**p<0.01, ***p<0.005, Kruskal Wallis test with Dunn’s post-hoc test for multiple comparison).

In the cortex, optogenetic activation of forebrain cholinergic input improves neuronal signal-to-noise ratio^26^. We used our results to predict how opposing cholinergic gain control alters this cholinergic computation by simulating how responses to weak (X_A_) and strong (X_B_) inputs would be transformed by the empirically measured IO functions of single CC and CS neurons (Fig 3a). Such analyses have been used to demonstrate the ability of norepinephrine to alter neuronal signal-to-noise ratio^27^. In CC neurons, for two inputs centred around 100 and 200 pA (with noise of 100 pA; Fig. 3a1), cholinergic action (Fig. 3a2) improved the separation of output distributions (Y_GA_, Y_GB_; Gain modulated output distributions) compared to basal conditions (Y_A_, Y_B_) by 56% (Fig 3a3, 3a4). However, for CS neurons the situation was reversed (Fig 3b): cholinergic input reduced the separation of output distributions by 35% (Fig 3b3, 3b4). This simulation indicates that cholinergic input improves the signal-to-noise ratio for CC neurons while reducing it for CS neurons; thus, co-release of ACh and GABA acts as a toggle to switch both the gain (Fig 2) and the signal-to-noise ratio of these projection neuron subpopulations. This effect occurs across a wide range of inputs for both CC and CS neurons (Fig S2a, S2b).

**Figure 3:**
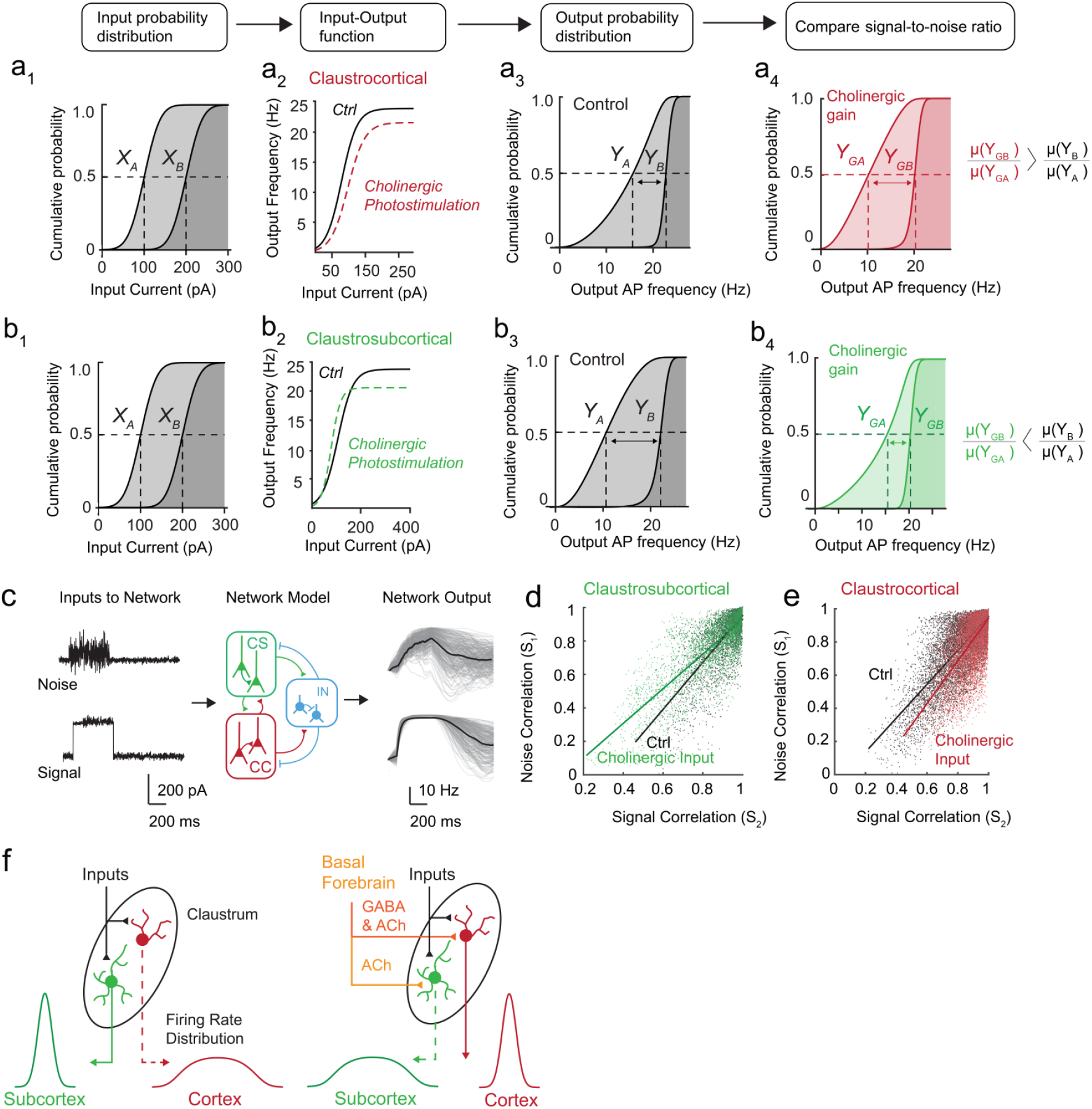
Incorporation of opposing gain control toggles signal-to-noise ratio and alters the correlation structure of model recurrent networks in a subpopulation specific manner. **a**. Paradigm used for signal-noise experiment: **a_1_**: Two inputs distributions (X_A_, X_B_) are transformed by empirical IO functions of CC (**a_2_**) to produce output distributions (Y_A_, Y_B_, **a_3_**). **a_4_**. Output distributions for CC neurons with IO functions with cholinergic photostimulation (Y_GA_, Y_GB_). **b**. The same inputs (**b_1_**) are transformed by the empirical IO functions of CS neurons (**b_2_**) to produce output distributions for with native IO functions (Y_A_, Y_B_, **b_3_**) compared to output distributions with IO functions with cholinergic photostimulation (Y_GA_, Y_GB_, **b_4_**) **c**: Paradigm used for modelling experiment: A signal (Gaussian distributed; μ = 200 pA, δ = 20 pA) and noise (Poisson distributed; μ = 100 pA, δ = 100 pA) are presented to an inhibition stabilized model of the claustrum. Right: Outputs of individual neurons (grey) and population average (black). **d**. Quantification of noise and signal correlations for CC neurons. (Linear fit, Ctrl: R^2^ = 0.55, Cholinergic input: R^2^ = 0.40) (B_2_). **e**. Quantification of noise and signal correlations for CS neurons. (Linear fit, Ctrl: R^2^ = 0.50, Cholinergic input: R^2^ = 0.57). **f.** Cell-type specific cholinergic gain modulation leads to a toggle between cortical and sub-cortical projections in the claustrum.

Recent studies demonstrate that cholinergic input improves network encoding efficiency by altering the relationship between signal and noise correlations in the neuronal activity of cortical networks^4,28^. A strong relationship between these is harmful for encoding a given signal, because it reduces discrimination of this signal from noise; in the cortex, cholinergic input - as well as attention - weakens this relationship^4,29,30^. We predicted the impact of cholinergic co-release on claustrum network correlation structure by using a recurrent circuit model based on an inhibition-stabilized network^31^. This model contained 300 neurons, including excitatory CC and CS projection neurons and the three inhibitory interneuron types; the IO function of each neuron type was defined by experimental measurements (See Methods, Fig S3). This network was driven by two types of stimuli similar to those previously applied *in vivo*^4^: (1) a signal with Gaussian amplitude distribution, with a SD of 1/10^th^ of the mean; and (2) Poisson distributed noisy input with a SD is equal to its mean (Fig 3c). For both stimuli, we examined the impact of cholinergic input on signal correlations and noise correlations between neurons. Remarkably, changing neuronal gain, by replacing native IO functions with IO functions measured during cholinergic photostimulation, decreased the slope of the signal correlation–noise correlation plot for CS neurons across a range of signal input sizes above 140 pA (*ΔSlope =* −15%, Fig 3d, Fig S2c), thus weakening the relationship between signal and noise correlations, as predicted from theory and *in vivo* experiments^4,32^. Because weakening this relationship leads to better signal discrimination, the reduction in correlation slope is associated with greater encoding capacity in networks^4,29,32^. In contrast, for CC neurons the slope increased for a range of signal input sizes above 140 pA^4^ *(ΔSlope =* +16%, Fig 3e, Fig S2d), thus strengthening the relationship between signal and noise correlations and reducing the encoding capacity of this population. Hence, we also observe a toggle in encoding efficiency between the CC and CS populations.

For signal inputs smaller than 100 pA, we observed a toggle in the opposite direction, with efficiency increasing for CC neurons while decreasing for CS neurons (Fig S2c, S2d). Our model indicates that the opposing cholinergic effects on gain due to co-release of ACh and GABA toggles network encoding efficiency in an input-dependent manner between distinct subpopulations of projection neurons, increasing efficiency for one population while reducing it for the other (Fig 3f).

Cholinergic modulation has been investigated in diverse experimental paradigms. Our results connect these observations and provide a microcircuit basis for the cholinergic control of signal-to-noise ratio and encoding capacity, based on opposing gain control of specific cell types in the claustrum. Our results highlight that cholinergic modulation does not affect networks uniformly: instead, it toggles information between subpopulations, from a cortically projecting to a subcortically projecting population in an input-dependent manner in the case of the claustrum (Fig 3f). This mechanism might also explain the ability of the claustrum to inhibit the cortex during slow-wave sleep^13^, where a low cholinergic tone would switch the discriminability of input towards the cortically-projecting claustral population. The ability to switch information between subpopulations might constitute a network mechanism to implement cholinergic control of attention^5^.

## Acknowledgments

We thank K.L.L Wong, Z. Chia, G. X. Ham, L. Mark, and G. Silberberg for insightful discussions and comments on our paper and K. Chung, P. Teo, S. Kay, R. Tan, and Y. C. Teo for technical assistance.

## Supplementary Figure Legends

**Supplementary Figure 1:**
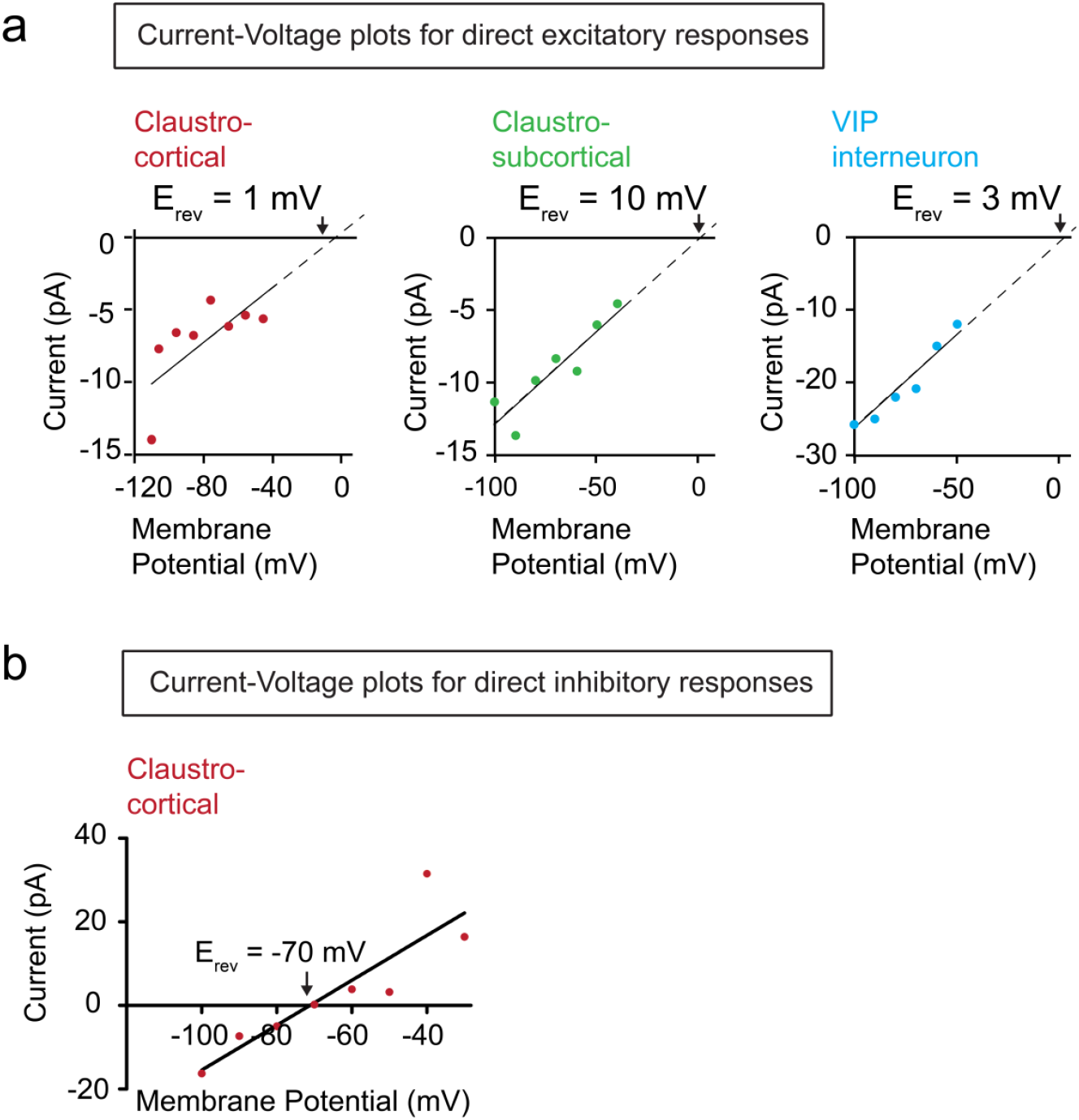
Current-Voltage relationship for direct cholinergic inputs. The amplitude of voltage clamp responses is measured for a range of holding potentials to determine the reversal potential of the response **a**. Current-Voltage plots for direct excitatory input for CC, CS and putative VIP neurons. Direct excitatory inputs are nicotinic receptor mediated as Erev is closest to the reversal potential of the nicotinic receptor (0 mV). **b**. Current-Voltage plots for direct inhibitory inputs reveals Erev closest to the reversal potential of chloride in internal solution (−80 mV).

**Supplementary Figure 2:**
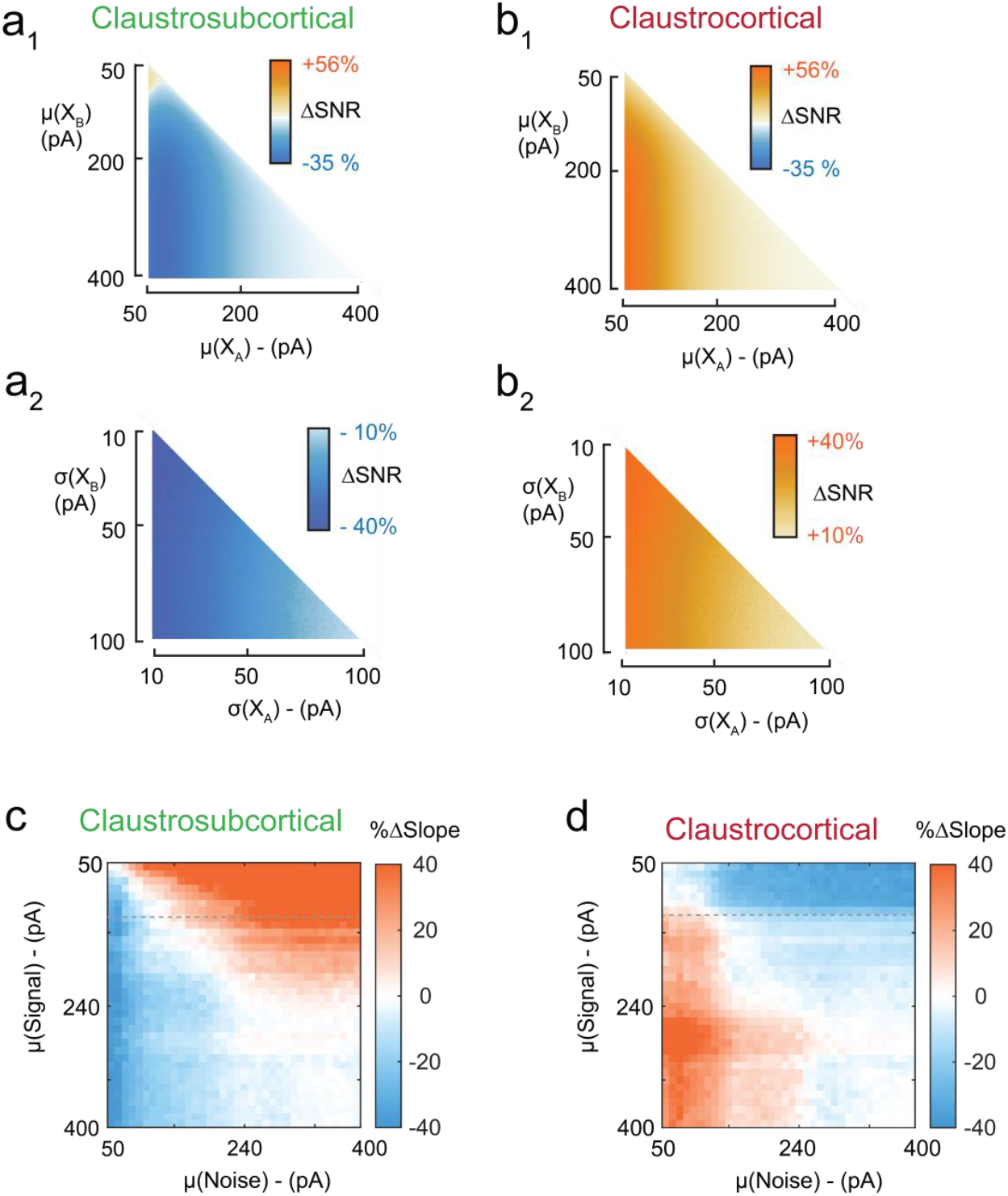
Examining signal-to-noise ratio and encoding efficiency for a range of inputs. **a**. Effect of varying input size (**a_1_**) and input noise (**a_2_**) of distributions X_A_ and X_B_ in Fig 3 on SNR for CS neurons. **b**. Effect of varying input size (**b_1_**) and input noise (**b_2_**) of distributions X_A_ and X_B_ on SNR for CC neurons. **c, d**. Dependence of correlations’ slope on input size, i.e mean of Gaussian-signal and mean of Poisson-noise for CS neurons (**c**) and CC neurons (**d**). The grey dotted line shows that the toggle is dependent on signal-input size and takes place in an opposite direction above 140 pA.

**Supplementary Figure 3:**
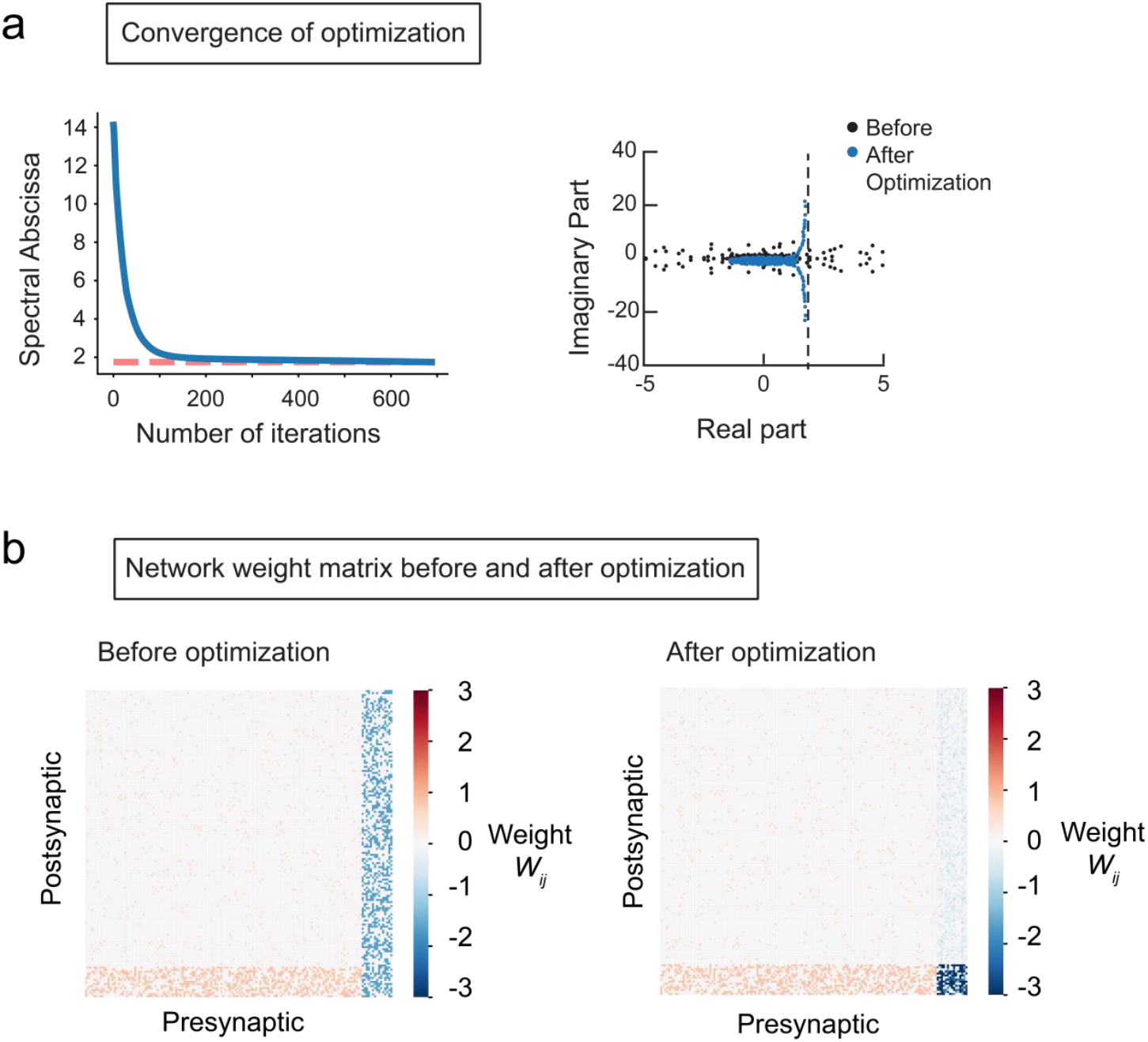
Optimization of inhibition stabilized claustrum model. The excitatory connectivity is specific by in-vitro experiments and previous findings^18^ while inhibitory weights are modified by removing the unstable eigenvalues of the weight matrix **W** towards stability^33^. **a**. Optimization of the model by refining the spectral abscissa (largest real part of the eigenvalues in **W**) over multiple iterations. **b**. Network weight matrix **W** before and after optimization process for all 300 neurons.

## Methods

### Animals

All animal experiments were performed according to the Guidelines of the Institutional Animal Care and Use Committee of Nanyang Technological University, Singapore (Protocol number: 151075). 35 adult ChAT-Cre x floxed ChR2-YFP (B6;129S6-*Chat^tm2(cre)Lowl^* /J; # 006410) mice of both sexes were used to study cholinergic input to CLA cells. The average age of mice used in our experiments was postnatal day 65 ±0.6.

### Brain slice recording

Acute brain slices were prepared according to the general procedures described in Graf et al. (2020). Mice were deeply anesthetized with isoflurane and euthanized via decapitation. The brains were isolated and transferred into ice-cold sucrose solution containing the following: 250 mM sucrose, 26 mM NaHCO_3_, 10 mM glucose, 4 mM MgCl_2_, 3 mM myo-inositol, 2.5 mM KCl, 2 mM sodium pyruvate, 1.25 mM NaH_2_PO_4_, 0.5 mm ascorbic acid, 0.1 mM CaCl_2_, and 1 mM kynurenic acid, with an osmolality of 350–360 mOsm and a pH of 7.4. Coronal brain slices (250 μm) were cut with a Leica VT 1000S vibratome. Slices were kept for 0.5 h at 34°C in artificial CSF (ACSF) containing the following: 126 mM NaCl, 24 mM NaHCO_3_, 1 mM NaH_2_PO_4_, 2.5 mM KCl, 2 mM CaCl_2_, 2 mM MgCl_2_, 10 mM glucose, and 0.4 mM ascorbic acid; 300–310 mOsm, pH 7.4, and gassed with a 95% O_2_/5% CO_2_ mixture before transfer to ACSF at room temperature for recordings.

Whole-cell patch clamp recordings were performed using Borosilicate glass pipettes (5-9 MΩ) filled with internal solution containing the following: 130 mM K-gluconate, 10 mM KOH, 2.5 mM MgCl_2_, 10 mM HEPES, 4 mM Na_2_ATP, 0.4 mM Na_3_GTP, 5 mM EGTA, 5 mM, Na_2_ phosphocreatinin, and 0.2% neurobiotin (290–295 mOsm, pH 7.4). Recordings were performed at 24°C with a MultiClamp 700B amplifier (Molecular Devices) and a Digidata 1440 interface (Molecular Devices). Signals were acquired at 50 kHz and filtered at 10 kHz. Access resistance (R_a_) was measured and only cells with R_a_ < 30 MΩ were used for further analysis. Cell-type identity was determined using an automated classifier using electrical properties described in Graf et al., 2020. Fourteen electrophysiological properties described in Graf et al., 2020 are extracted using software made available by the authors at https://github.com/adityanairneuro/claustrum. A trained classifier is used to distinguish between the two subtypes of claustral projection neurons and three subtypes of claustral interneurons.

For optogenetic photoactivation of cholinergic terminals, slices were illuminated by a 130W short arc mercury lamp (Olympus) passed through an EYFP filter set and a x25 water-immersion objective. For voltage clamp experiments, we delivered 50 ms light pulses while clamping neurons at −40 mV. For current clamp experiments, we delivered blue light at 10Hz in 20 ms light pulses for a duration of one second. We choose this stimulation protocol to mimic average firing rates of basal forebrain cholinergic neurons which range from 7-14 Hz during wakefulness and REM sleep^34,35^.

### Analysis of neuronal gain

In current clamp experiments, we constructed input-output (IO) curves for each neuron by injecting depolarizing current pulses in the range 0-400 pA in 20 pA steps and measuring output firing frequency. Empirically determined input current-output frequency curves were fit with sigmoidal tanh functions of the form:

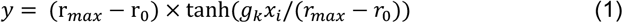

Where *r_max_* is the maximum observed firing frequency, *r_0_* is the baseline firing frequency, *x* is the input current and *g_k_* is the gain or slope of the function at baseline and thus represents the input-output sensitivity of the neuron *k*.

### Analysis of signal-to-noise ratio with simulations using empirical IO functions

Given that cholinergic input can alter gain, we analyzed whether these empirically observed differences in IO curves of CC and CS neurons might be sufficient for these projections to process input differently. We modelled this possibility by considering how two input probability distribution functions (PDF: X_A_, X_B_) would be transformed by IO functions of neurons with and without cholinergic photostimulation (Fig 3a). To quantify the amount of separation between output PDFs, we compared the change in the ratio of their means with and without cholinergic gain control as this reflects the SNR of the two signals X_A_ and X_B_^27^. Formally:

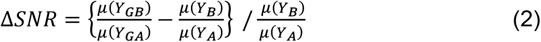

Where Y_GB_ and Y_GA_ are output PDFs in the presence of cholinergic gain control whereas Y_B_ and Y_A_ are outputs PDFs in its absence. *μ(Y)* indicates the mean of distribution Y.

We verified that the SNR results we observed generalized for a range of input (PDF: X_A_, X_B_) means and SDs by systematically varying either the mean of X_A_, X_B_ (Fig S2a_1_, S2b_1_) or the SD of X_A_, X_B_ (Fig S2a_2_, S2b_2_).

### Analysis of signal and noise correlations in model CLA-like networks

To understand the role of cholinergic gain control in CLA-like networks, we constructed a recurrently connected network model using stability optimised circuits (SOCs), a class of networks where inhibition stabilises the network to create a non-chaotic network with transient dynamics. Below we briefly describe this model.

We first generate synaptic weight matrices ***W*** with *N* = 300 neurons (with excitatory and inhibitory neurons in the ratio 9:1 as empirically determined^36^) as detailed in Hennequin et al., 2014^31^.

We begin with set of sparse weights with non-zero elements set to 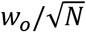 for excitatory neurons and 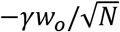 for inhibitory neurons, where 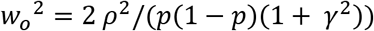 with connection probability p being 0.03 for excitatory neurons and 0.4 for interneurons as empirically determined^37^. We construct ***W*** with an approximately circular spectrum (i.e set of eigenvalues) of radius ρ = 10 and inhibition/excitation ratio γ = 3 in line with Hennequin et al., 2014 (Fig S3b, Left).

Following construction of ***W***, we never change the excitatory weight, but refine the inhibitory connections to minimize the ‘spectral abscissa’ of W, which is the largest real part among the eigenvalues of W (Fig S3a). This optimization is performed according to Stroud et al., 2018 and the resulting matrix, referred to as a stability optimized circuit or SOC is non-chaotic^38^ (Fig S3b, Right).

The use of SOCs is an approximation used due to the lack of precise cell-type specific connectivity for the CLA. SOCs have been used to study the effect of gain modulation in motor cortex circuits^38^. Since we were interested in gain control of the CLA network, we used SOCs to obtain a non-chaotic network with CLA-like connectivity for PN-PN and IN-IN connections and empirically determined IO curves for each cell type.

Our model is governed by a differential equation which controls neuronal activity (Equation 3) using the gain function (Equation 1) and the synaptic connectivity matrix ***W***.

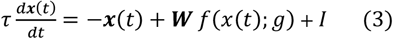

We integrate Equation 1 using the ODE45 function in Matlab using default parameters.

The initial condition *x_0_* was chosen among the ‘’most observable’ modes which elicit the strongest transient dynamics according to Hennequin et al., 2014^31^.

To mimic in-vivo experiments which examined the effect of cholinergic input on signal and noise correlations in networks^39^, we delivered time varying inputs *I* as shown in Fig 3 C. We delivered two sets of inputs, a signal which consists of gaussian distributed input where the SD of input is 1/10^th^ of its mean. In a different set of trials, we provided a second noisy input to the network which was Poisson distributed with SD equal to the mean of the signal. Signal correlation – Noise correlation graphs are obtained by plotting the Pearson’s correlations coefficient (PCC) between every pair of excitatory neurons during the presentation of the Gaussian signal vs during the presentation of Poisson noise.

To ensure that the results we observed were robust for a range of input sizes, we systematically varied the mean of either the Gaussian signal or Poisson noise and examined the slope of the signal-noise correlation plot (Fig S2 c, S2d).

Code used for analysis of IO curves and the recurrent claustrum network is available at https://github.com/adityanairneuro/cholinergic

